# The phylogenetic signal in primate ontogenies, with special attention to dental development

**DOI:** 10.64898/2026.03.12.710081

**Authors:** Paola Cerrito

## Abstract

Comparative studies in evolutionary biology must account for trait non-independence arising from shared ancestry. While the phylogenetic signal of adult traits has been extensively studied, little is known about how conserved developmental trajectories are across species. Here, I quantify the phylogenetic signal (K) in the ontogeny of 35 traits across 157 primate species, spanning motor, cognitive, life-history, and dental development. Using Blomberg’s K statistic and a species-level mammalian phylogeny, I test two predictions: (i) that morphological (dental) traits exhibit the strongest phylogenetic signal, and (ii) that earlier-developing traits are more conserved. Results show that life-history traits are the most phylogenetically labile, while dental development is the most conserved (K = 0.7-2.6), with the eruption of the mandibular canine showing the highest signal (K = 2.6). Contrary to expectations, later-developing traits, particularly permanent teeth, display stronger phylogenetic conservation than earlier-developing deciduous teeth. These findings suggest that even within a single developmental system, the strength of phylogenetic constraint varies markedly with timing. The results provide an empirical foundation for identifying reliable temporal anchors in comparative primate ontogeny and have implications for interpreting maturational patterns in human evolution and the fossil record.

## Introduction

Comparative analyses must deal with trait non-independence, meaning that closely related species tend to resemble each other. A central question in many evolutionary studies, is understanding how much such resemblance is a consequence of shared ancestry, as opposed to being the result of similar adaptive strategies or selective pressures. A widely used measure of phylogenetic signal is the descriptive statistic *K*, originally published by Blomberg and colleagues (2003). They analyzed 121 different traits, and found that “behavioral traits exhibit lower signal than body size, morphological, life-history, or physiological traits”. Similar results have been found when focusing the analyses on primates (Kamilar & Cooper, 2013). However, these results pertain to the adult phenotype. Conversely, little is known regarding the phylogenetic signal of ontogenetic data. That is: which traits are more conserved than others in the age at which they develop? This question, despite being largely unanswered, is in my opinion central in evo-devo studies, which account for ontogenetic trajectories in explaining the adult phenotype.

For example, assessing which traits have undergone a recent change in their ontogenetic trajectory is essential to many studies on human evolution. Yet, to assert that a given developmental milestone occurs “earlier” or “later” than expected, one must first define in relation to what it is “earlier” or “later”. Essentially one must confirm that it is the trait in question, and not the temporal marker against which one assesses it, that has undergone evolutionary changes in its timing. One can imagine sitting in train A and seeing that it is moving relative to train B that is next to it: are both trains moving, is only A or only B moving? A classic example is the relationship between age at weaning, brain maturation and dental development in primates. Starting with Schultz’s seminal work (Schultz, 1969), a large amount of research (e.g. Godfrey et al., 2001; T. M. Smith, 2013, 2013) has been devoted to understanding the relative temporal relationships between dental development, brain ontogenies and life histories. Moreover, many studies on human evolution use ages of dental development for proxies as changes in developmental trajectories more broadly (R. J. Smith et al., 1995; T. M. Smith et al., 2010; T. M. Smith, Tafforeau, et al., 2007; T. M. Smith, Toussaint, et al., 2007). Yet, the question of what is accelerated in relation to what, and what is delayed in relation to what remains largely unresolved. All variables (dental development, weaning, sexual maturation, brain growth, etc.) can be moved relative to each other along the axis of time, failing to be rooted by something that has been shown to serve as a good anchor. Furthermore, it also not understood which variables (traits) are more constrained in their timing and which are more amenable through change during evolutionary times, thus potentially being indicative of directional change rather than phylogenetic history.

While it might seem intuitive to elect birth as time zero (t_0_) against which to measure all other ages at event occurrence, data showing that gestation length is phylogenetically conserved would be necessary to substantiate this assumption, and one might suspect that it is not, as it has been shown that life history traits are quite labile (Blomberg et al., 2003). Similarly, comparing the duration of ontogenetic stages by operationalizing them as proportion of lifespan, is also a discussible choice since lifespan is another very labile trait (Blomberg et al., 2003), and only recently a few studies have begun to use age from conception as temporal anchor (Charvet et al., 2023; Gómez-Robles et al., 2024).

Here, I explicitly address this issue by measuring the phylogenetic signal of a variety of traits, and types of traits, in primates. My aim is to provide a concrete and practical tool for primate evo-devo studies – which are especially relevant since so much that is considered to be unique about them, such as tool use (Heldstab et al., 2020), socio-cognitive development (Cerrito et al., 2024) and prosocial behaviors (Hrdy et al., 2022) has to do with the way in which they grow and develop. Specifically, I aim to assess the strength of the phylogenetic signal in different categories of variables, such as motor, cognitive, dental and physiological development, in order to understand which ones are constrained and may serve as a good temporal anchor in comparative studies, and which ones are labile and more likely subject to change in repones to evolutionary forces (be it mutation, selection, drift or gene flow). I make the following the predictions: i) based on previous findings on the adult phenotype (Blomberg et al., 2003; Kamilar & Cooper, 2013), I predict that phylogenetic signal is the strongest in the ontogeny of morphological traits (specifically: dental); ii) since later developing traits can be more subject to environmental influences, I predict that a lower average age at trait emergence is correlated with a stronger phylogenetic signal.

## Materials and methods

### Data collection

I collected from the literature (Abondano et al., 2022; Ausderau et al., 2017; Brown & Dixson, 2000; Byrne & Suomi, 1995; Carruth & Skinner, 2002; Cerrito & Spear, 2022; Chalmers, 1980a, 1980b; Chism, 1986; Cockrum, 1962; Doran, 1997; Dunbar & Badam, 1998; Eaglen & Boskoff, 1978; Ehrlich, 1974; Ferrari, 1992; Garcia et al., 2006; Gauthier, 1999; Gould, 1990; Govindarajulu et al., 1993; Guthrie & Frost, 2011; Heldstab et al., 2020; Hoff et al., 1981; Jogahara & Natori, 2012; King et al., 1974; Kinoshita et al., 2017; Klopfer & Klopfer, 1970; Lee et al., 2020; Malalaharivony et al., 2021; Mittermeier et al., 2022; Nash, 2003; Peng et al., 1973; Potì & Spinozzi, 1994; Rademacher, 2024; Rosenson, 1972; Setchell & Wickings, 2004; Sievert et al., 1991; B. H. Smith et al., 1994; Tacutu et al., 2012; Vogt et al., 1978; Wang et al., 2014; Young & Shapiro, 2018; Zimmermann, 1989) data of age at event occurrence (in days, starting from conception) for 35 variables (Table 1) in four different categories (motor development, dental development, cognitive development, life history). The average sample size per variable is 39 species. The total number of primate species represented is 157. When data was available for both males and females, the average value was taken. The complete dataset, together with sources for each datapoint is available as supplementary material (Data S1). Since most sources express their values in age after birth, following Charvet et al. (2023) I added gestation duration to obtain age since conception.

**Table 1.** | For each of the traits analyzed: Category; phylogenetic signal (K); significance of K; number of species (sample size) and average age, among species at which the trait occurs / emerges (expressed in days, log-transformed).

| Category | Variable | K | p.value | Number of species | Log (Mean age (days)) |
| --- | --- | --- | --- | --- | --- |
| Cognitive | Onset of social play | 0.77 | 0.00918 | 14 | 5.44 |
| Dental Development | Eruption C Mandible | 2.64 | 0.00001 | 20 | 7.01 |
| Dental Development | Eruption C Maxilla | 2.45 | 0.00001 | 21 | 7.12 |
| Dental Development | Eruption Deciduous C Mandible | 1.8 | 0.00001 | 25 | 5.44 |
| Dental Development | Eruption Deciduous C Maxilla | 1.39 | 0.00001 | 25 | 5.41 |
| Dental Development | Eruption Deciduous I1 Mandible | 1.14 | 0.00001 | 25 | 5.22 |
| Dental Development | Eruption Deciduous I2 Mandible | 0.97 | 0.00001 | 25 | 5.26 |
| Dental Development | Eruption Deciduous I2 Maxilla | 0.92 | 0.00001 | 25 | 5.3 |
| Dental Development | Eruption Deciduous P3 Mandible | 1.21 | 0.00001 | 25 | 5.4 |
| Dental Development | Eruption Deciduous P3 Maxilla | 0.89 | 0.00001 | 25 | 5.42 |
| Dental Development | Eruption Deciduous P4 Mandible | 1.45 | 0.00001 | 25 | 5.57 |
| Dental Development | Eruption Deciduous P4 Maxilla | 0.99 | 0.00001 | 25 | 5.67 |
| Dental Development | Eruption I1 Mandible | 2.17 | 0.00001 | 21 | 6.67 |
| Dental Development | Eruption I1 Maxilla | 1.97 | 0.00001 | 20 | 6.81 |
| Dental Development | Eruption I2 Mandible | 2.47 | 0.00001 | 20 | 6.75 |
| Dental Development | Eruption I2 Maxilla | 2.09 | 0.00001 | 19 | 6.92 |
| Dental Development | Eruption M1 Mandible | 1.18 | 0.00001 | 25 | 6.18 |
| Dental Development | Eruption M1 Maxilla | 1.59 | 0.00001 | 25 | 6.27 |
| Dental Development | Eruption M2 Mandible | 1.88 | 0.00001 | 25 | 6.68 |
| Dental Development | Eruption M2 Maxilla | 1.91 | 0.00001 | 24 | 6.71 |
| Dental Development | Eruption M3 Mandible | 1.9 | 0.00001 | 23 | 7.2 |
| Dental Development | Eruption M3 Maxilla | 2.39 | 0.00001 | 22 | 7.24 |
| Dental Development | Eruption P3 Mandible | 1.71 | 0.00001 | 22 | 7.01 |
| Dental Development | Eruption P3 Maxilla | 2.29 | 0.00001 | 21 | 6.98 |
| Dental Development | Eruption P4 Mandible | 1.59 | 0.00001 | 19 | 7.11 |
| Dental Development | Eruption P4 Maxilla | 2.04 | 0.00001 | 18 | 7.1 |
| Dental Development | Eruption Deciduous I1 Maxilla | 0.7 | 0.00004 | 25 | 5.28 |
| Life History | Age at sexual maturity | 1.01 | 0.00001 | 25 | 6.88 |
| Life History | Weaning | 0.97 | 0.00001 | 25 | 6.04 |
| Life History | Age at first reproduction | 0.95 | 0.00003 | 25 | 7.37 |
| Life History | Gestation duration | 0.6 | 0.00021 | 25 | 5.08 |
| Life History | Maximum longevity | 0.51 | 0.01611 | 25 | 9.17 |
| Motor | Locomotor independence | 1.15 | 0.00001 | 25 | 6.08 |
| Motor | Adult level skill competence reached | 1.15 | 0.00002 | 25 | 5.71 |
| Motor | Onset of crawling | 1.03 | 0.00045 | 14 | 5.38 |

### Data analysis

All data is expressed in days and has been log-transformed before analyses. I measured the phylogenetic signal of each variable using K (Blomberg et al., 2003). The analyses used the species-level mammal phylogeny from Upham and colleagues (Upham et al., 2019). I used their maximum clade credibility tree calibrated using node dates and an exponential prior. To determine the significance of K, the measured value of K is compared against a distribution of K values generated by iteratively randomly redistributing the data across the phylogeny. I used the phylosig function in the “phytools” package (Revell, 2012) in R to calculate K and I determined its significance by running 100,000 iterations.

Given the uneven distribution of the sample in terms of number of species present per variable analyzed (which ranges from 14 for Onset of crawling to 157 for Gestation duration) I proceeded to first assess the effect of sample size on phylogenetic signal. I chose the eight variables for which the sample size was much larger than for others (gestational age at birth, n=157; maximum age at death, n=136; age at weaning, n=117; age at first reproduction, n=113; age at sexual maturity, n=135; age at locomotor independence, n=36; age at which adult-level manual skill competence is reached, n=32; age at mandibular M1 eruption, n=32), iteratively randomly selected 11 species 500 times, and proceeded to compute the phylogenetic signal (K). I repeated the same operation capping the sample size at 20 instead of 11. I then calculated the average value and standard deviation of K over the 500 iterations. Then, based on the results obtained above and on previous research (Blomberg et al., 2003), I proceeded to calculate K for all the variables present in the dataset, capping the sample size at 25.

Subsequently, to assess the relationship between phylogenetic signal and age at trait emergence I focused on dental development, which contains 26 different traits of a same category and therefore minimizes inter-categorical difference as a source of variance. I constructed seven linear models with K as response variable and: i) average age as predictor; ii) average age and tooth type (permanent or deciduous) (as additive term) as predictors; iii) average age and tooth type (permanent or deciduous) (as interaction term) as predictors; iv) average age and jaw (maxilla or mandible) (as additive term) as predictors; vi) average age and jaw (maxilla or mandible) (as interaction term) as predictors; vii) average age and sample size (as additive term) as predictors; v) average age and sample size (as interaction term) as predictors. I then compared the seven models using Akaike’s information criterion (AIC) (Akaike, 1973) and selected as best ones the one with an AIC value within two of the lowest one.

Finally, to better understand whether variation in K across traits may be produced by different ways of departing from Brownian motion expectations I proceeded with fitting different evolutionary models to the traits and then comparing them using AIC. Specifically, to each trait I fitted these six different models: Brownian Motion (BM) (Felsenstein, 1985), Ornstein–Uhlenbeck (OU) (Hansen, 1997), Early Burst (EB) (Harmon et al., 2010), lambda: λ (Pagel, 1999), delta: δ (Pagel, 1999), and kappa: κ (Pagel, 1994). To fit the different models I used the fitContinous function in the R package “geiger” (Pennell et al., 2014). OU assumes that traits are pulled towards and optimal value, that is: stabiliaing selection toward an optimum. EB assumes a rapid evolution of the trait early in a clade’s history (for example during an adaptive radiation) and then a slow doen over time. BM assumes that evolutionary changes are random and accumularte orver time, hence differences in species are proportional to their phylogenetic history. Lambda (λ), similarly to Blomberg’s K, essentailly measures phylogenetic signal. Delta is a time-dependent transformation which is designed to caputre variation in rates of evolution through time, with all elements of the variance-covariance matrix raised to a given power (δ). Finally, kappa, operates a transformation on branch lengths, and can be understood as a model in which change is more or less concentrated at speciation events. All models estimated an evolutionary rate parameter (σ²), with some models additionally estimating parameters describing stabilizing selection (OU: α), rate variation through time (EB: a; δ), or phylogenetic signal scaling (λ, κ). Model support was assessed using AIC, which penalizes model complexity appropriately.

## Results

As reported in Table S1, phylogenetic signal tends to decrease with increasing sample size. Furthermore, and consistently with previous work (Blomberg et al., 2003), with sample sizes of 11 K was mostly not significant (at an alpha threshold of 0.05).

Overall, the most phylogenetically liable traits are life-history ones, while the most conserved ones are those relative to dental ontogeny. However, the ranges within each category greatly overlap between each other, such that K is between 0.7 and 2.6 for dental traits; between 0.6 and 1 for life history traits; between 1 and 1.15 for motor development. The complete results are visible in Figure 1 and reported in Table 1. The most labile trait is maximum longevity (K = 0.51), while the least labile is age at eruption of the mandibular canine (K = 2.5). These results mostly support my first prediction: that dental ontogeny is more conserved than physiological (life history traits) or behavioral one (motor and cognitive traits).

**Figure 1.**
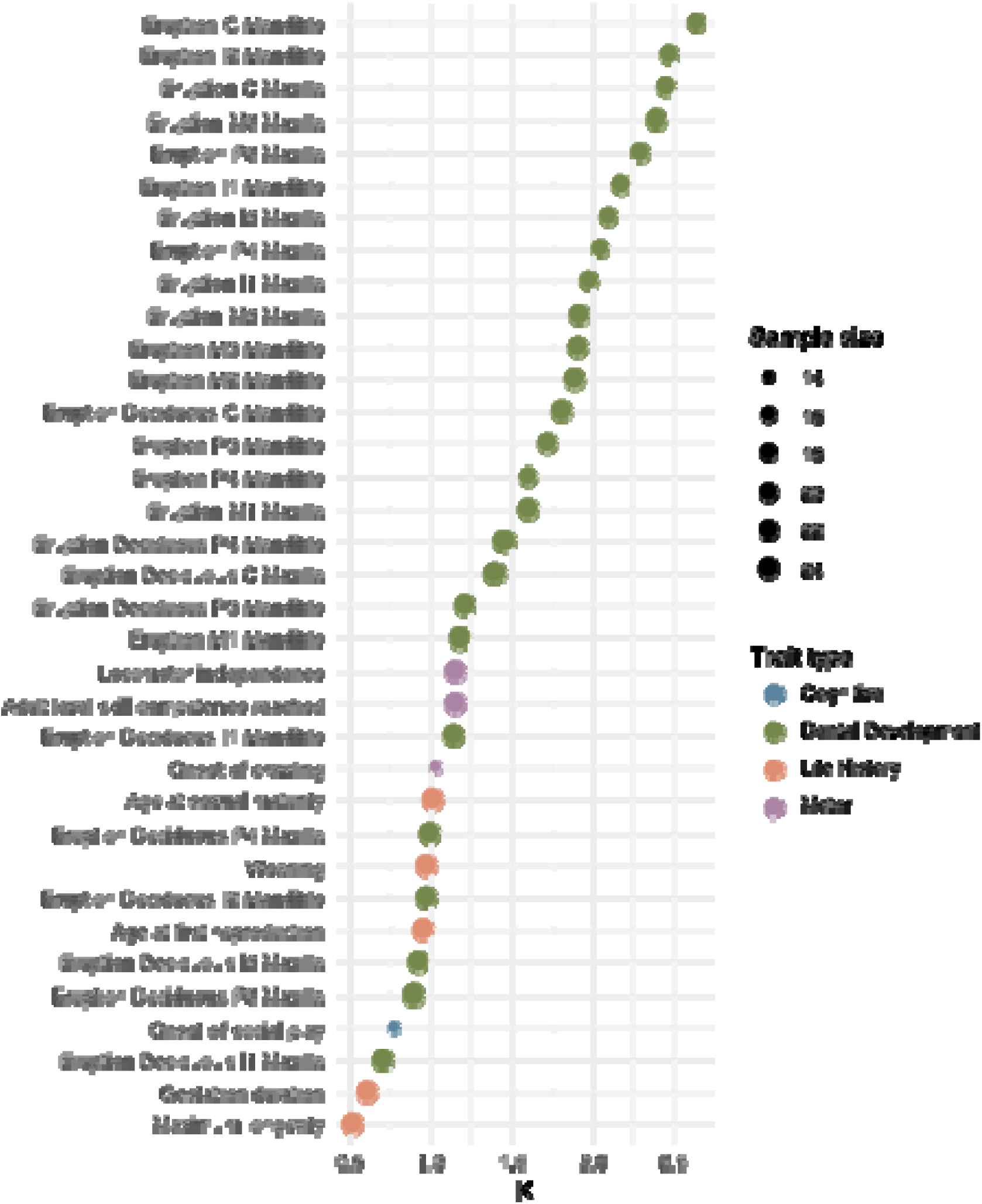
| Phylogenetic signal (K) for each of the traits analyzed. I = incisor; C = canine; P = premolar; M = molar. Dot size corresponds to number of species sampled for each variable.

The results regarding the relationship between age at trait emergence and phylogenetic signal strongly contrast my second prediction: age at dental emergence seems to directly, rather than inversely, correlate with K. The direct correlation is weak when considering all traits together, but becomes very strong when separating them by category, and remains positive (slope = 0.6) for dental development while it becomes slightly negative (–0.05) for life history traits. These results can be visualized in Figure 2a. Moreover, the results for dental development indicate that permanent teeth have a stronger phylogenetic signal than deciduous ones (Figure 2b).

**Figure 2.**
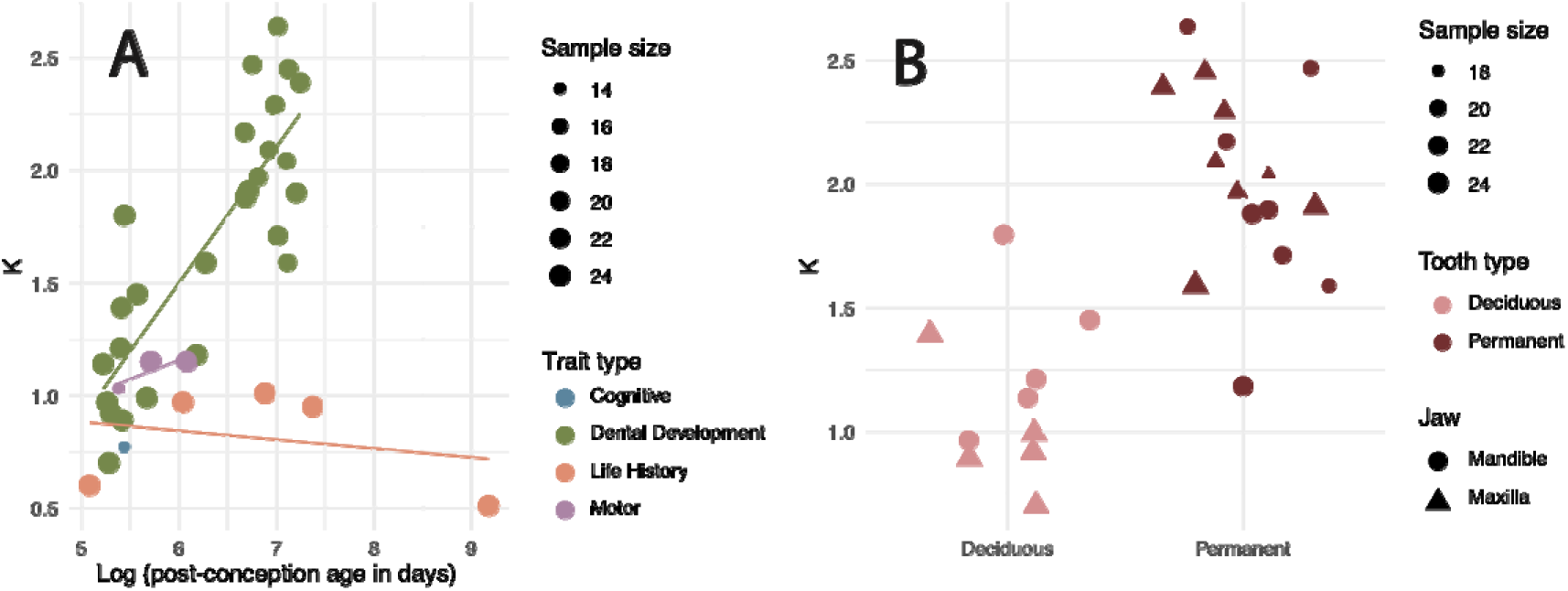
| **A**: For each of the traits analyzed, phylogenetic signal (K) is plotted against the mean age, in days, at which the trait occurs. Age values have been log-transformed for both the analyses and the figure. Dot color represents the category of the trait, dot size represents the number of species sampled for that trait. **B:** Phylogenetic signal of dental emergence shown per tooth type, permanent or deciduous.

The results of the comparisons between the seven models I constructed (Table 2) to explain the variance in K between ages at emergence of different teeth, indicate the age at emergence alone is one of the best models, hence it is not necessary to include either tooth type (permanent or deciduous) or jaw (mandible or maxilla) to explain the observed differences in phylogenetic signal.

**Table 2.** | For each of the models constructed: the adjusted R-squared value, the AIC score and a variable indicating whether the model in question is the best one (or within two of the best one).

| Model | R-squared | AIC | Best |
| --- | --- | --- | --- |
| $K \sim \text{Mean} * \text{Jaw}$ | 0.70 | 17.75 | yes |
| $K \sim \text{Mean}$ | 0.68 | 18.06 | yes |
| $K \sim \text{Mean} * n$ | 0.70 | 18.29 | yes |
| $K \sim \text{Mean} + n$ | 0.68 | 19.05 | yes |
| $K \sim \text{Mean} + \text{Jaw}$ | 0.67 | 19.70 | yes |
| $K \sim \text{Mean} + \text{Tooth\_type}$ | 0.67 | 19.90 | no |
| $K \sim \text{Mean} * \text{Tooth\_type}$ | 0.65 | 21.90 | no |

Model comparison results for all traits are summarized in Table S2. For the majority of traits, Brownian motion (BM) was included among the best-supported models (ΔAIC ≤ 2), indicating that trait evolution is generally well described by gradual divergence along the phylogeny. In contrast, BM was not supported for six traits, all of which are related to life-history timing, with the exception of one motor trait. Specifically, maximum longevity, gestation duration, age at weaning, age at sexual maturity, age at first reproduction, and age at locomotor independence were best explained by rate-scaling models, including Pagel’s λ and/or κ. For these traits, λ estimates were consistently high (λ ≈ 0.96–0.97), and κ estimates were substantially below one (κ ≈ 0.22–0.59), indicating strong phylogenetic structure combined with heterogeneous evolutionary rates. No trait was best supported by Ornstein–Uhlenbeck or early-burst models alone.

## Discussion

Although Brownian motion (BM) was included among the best-supported models for most traits, six traits showed clear departures from BM expectations (Table S2): maximum longevity, gestation duration, age at weaning, age at sexual maturity, age at first reproduction, and age at locomotor independence. These traits were best explained by rate-scaling models (Pagel’s λ and/or κ), indicating that their evolution is structured by phylogeny but does not follow a simple pattern of gradual divergence. Importantly, these departures from BM occur despite substantial variation in phylogenetic signal. For several of these traits, Blomberg’s K was relatively low to moderate (K = 0.51–0.95 for longevity, gestation duration, weaning, and age at first reproduction), indicating weaker resemblance among close relatives than expected under BM. At the same time, these traits showed high estimates of λ (λ ∼ 0.96–0.97), suggesting that trait covariation remains strongly structured by the phylogeny. This combination implies that evolutionary changes are phylogenetically patterned but unevenly distributed across the tree, rather than accumulating gradually within clades. Support for Kappa (κ) models, with κ values substantially below one (κ ∼ 0.22–0.59), further indicates heterogeneous evolutionary rates, with changes concentrated on shorter branches or early in lineage divergence. This pattern is particularly pronounced for age at sexual maturity, which showed exclusive support for κ and a very low κ estimate (κ ∼ 0.22). Such episodic evolution is consistent with rapid shifts associated with transitions between life-history strategies, followed by relative stasis within lineages. In contrast, age at locomotor independence showed higher phylogenetic signal (K = 1.15), consistent with stronger conservatism among close relatives. Nonetheless, this trait also departed from BM expectations, indicating that high phylogenetic signal does not necessarily imply gradual Brownian evolution. Together, these results demonstrate that variation in phylogenetic signal among ontogenetic traits reflects differences in how evolutionary change is distributed across the phylogeny, rather than differences in the overall strength of phylogenetic constraint.

The results of the analyses seem to confirm that traits that are more phylogenetically labile in the adult phenotype, are also more labile in their ontogenetic trajectories. However, the difference in ranges between categories that I report here, could derive from uneven sample sizes: there is one variable for cognitive development, three for motor development, five for life history traits and 30 for dental development. Furthermore, while traits relating to physiological development (life history) tend to have a lower phylogenetic signal than those related to skeletal development, the results of my analyses (Table S1) indicate that this is not an effect of an average larger sample size for life history traits than for the others as this remains true when capping the sample size of life history variables at a number comparable to those of skeletal traits. These results also confirm previous findings (Blomberg et al., 2003), indicating that significance is usually reached with 20 species.

While the over-representation of dental development traits may hinder the validity of between-category comparisons in the phylogenetic signal of the ontogeny of traits; it nonetheless provides a few very interesting results. First, it demonstrates that even within a given category of trait there is huge variance in phylogenetic signal (from 0.7 to 2.6). Second, it allows for a robust intra-categorical analysis of the relationship between K and age at trait emergence. This analysis highlights (Figure 2) that, contrary to expectations, later-developing teeth are much more conserved in the timing of their emergence than earlier-developing ones, and more generally, that the timing of permanent teeth emergence is much more conserved than the timing of deciduous teeth emergence. This finding alone has important implications in comparative studies of primates and humans, since the age at emergence of the first permanent molar (M1) is frequently taken as the temporal anchor against which to assess the earlier or later development of other features (Bermúdez de Castro et al., 2010; Godfrey et al., 2001; GuatellilJSteinberg, 2009; B. H. Smith, 1989; R. J. Smith et al., 1995). For example, it is frequently cited as evidence of humans’ early weaning, the fact that it occurs much earlier than M1 emergence (T. M. Smith, 2013). Interestingly, however, the results of my analyses indicate that M1 is far from being the most phylogenetically constrained tooth in the time of its emergence. Hence, other traits, such as mandibular canine emergence, might be best suited for these types of comparative developmental discussions.

Conversely, the age at emergence of some of the deciduous teeth have a rather low phylogenetic signal (K < 1). Given their low degree of constraint, they might constitute little explored (B. H. Smith, 2024) skeletal markers of labile maturational processes such as age at weaning or brain development, with important implications for human evolution and applications to the hominin fossil record, in which the timing of most developmental events has to be inferred from skeletal elements (T. M. Smith, Toussaint, et al., 2007; Zollikofer et al., 2024).

More broadly, these findings underscore the importance of a developmental perspective in studies of primate and human evolution. Evolutionary developmental (evo-devo) approaches depend critically on distinguishing between ontogenetic traits that are tightly constrained and those that are evolutionarily flexible. Traits characterized by strong phylogenetic conservation may reflect deep developmental integration and long-standing constraints, whereas more labile traits may be more responsive to ecological, life-history, or selective pressures. Identifying which aspects of ontogeny fall into each category is therefore essential for interpreting evolutionary change, especially when reconstructing developmental patterns in extinct taxa. By explicitly quantifying variation in phylogenetic signal across ontogenetic traits, this study provides a framework for identifying which developmental markers are most informative for evolutionary inference and which are most likely to capture adaptive or plastic responses in primate and human evolution.

## Data accessibility

all data is available will be available here as supplementary material upon publication: https://doi.org/10.6084/m9.figshare.31678663

